# Automatic identification and annotation of MYB gene family members in plants

**DOI:** 10.1101/2021.10.16.464636

**Authors:** Boas Pucker

## Abstract

**Background:** MYBs are among the largest transcription factor families in plants. Consequently, members of this family are involved in a plethora of processes including development and specialized metabolism. The MYB families of many plant species were investigated in the last two decades since the first investigation looked at *Arabidopsis thaliana*. This body of knowledge and characterized sequences provide the basis for the identification, classification, and functional annotation of candidate sequences in new genome and transcriptome assemblies.

**Results:** A pipeline for the automatic identification and functional annotation of MYBs in a given sequence data set was implemented in Python. MYB candidates are identified, screened for the presence of a MYB domain and other motifs, and finally placed in a phylogenetic context with well characterized sequences. In addition to technical benchmarking based on existing annotation, the transcriptome assembly of *Croton tiglium* and the annotated genome sequence of *Castanea crenata* were screened for MYBs. Results of both analyses are presented in this study to illustrate the potential of this application. The analysis of one species takes only a few minutes depending on the number of predicted sequences and the size of the MYB gene family. This pipeline, the required bait sequences, and reference sequences for a classification are freely available on github: https://github.com/bpucker/MYB_annotator.

**Conclusions:** This automatic annotation of the MYB gene family in novel assemblies makes genome-wide investigations consistent and paves the way for comparative studies in the future. Candidate genes for in-depth analyses are presented based on their orthology to previously characterized sequences which allows the functional annotation of the newly identified MYBs with high confidence. The identification of orthologs can also be harnessed to detect duplication and deletion events.

## Introduction

MYB transcription factors are named after an Avian myeloblastosis virus protein (v-Myb) which is a modified version of the cellular c-Myb and causes the activation of oncogenes [1]. While MYBs were first discovered in animals, they appear in substantially larger numbers in plants and form one of the largest transcription factor families [2–6]. A characteristic MYB feature is the presence of a conserved DNA-binding domain at the N-terminus [7]. Up to four imperfect amino acid repeats (50-53 amino acids) form three alpha-helices each [7]. Helix two and three of each repeat are arranged to a helix-turn-helix structure [8]. Three regularly spaced tryptophan or other hydrophobic amino acid residues form the core of this structure [8]. The third alpha helix is responsible for the direct DNA interaction [9]. The repeats are classified into R1, R2, and R3 based on similarity to the respective repeats of the first characterized MYB, c-Myb [1]. MYB proteins are classified based on the presence of these repeats. For example, R2R3-MYBs harbor the R2 and R3 repeat while 3R-MYBs have one copy of each of the repeats (R1R2R3). Further classification into subgroups can be achieved based on the phylogenetic relationships and characteristic sequence motifs in the C-terminal region [2, 5, 10]. Different MYB classification systems were proposed in previous studies [2, 5, 10].

R1R2R3-MYBs have been proposed to be regulators of the cell cycle with conserved functions between animals and plants [11, 12]. R2R3-MYBs account for the large MYB family size in plants [2]. The evolutionary origin of R2R3-MYBs and 3R-MYBs is still debated. The loss model proposes that R2R3-MYBs diverged from 3R-MYBs through loss of the R1 repeat [13–15], while the gain model proposes that the 3R-MYBs evolved from the R2R3-MYB through duplication of a repeat [10, 16]. R2R3-MYBs are involved in the regulation of numerous processes including the regulation of developmental processes, response to environmental stresses, and specialized metabolism [17–19]. WEREWOLF/MYB66 is a negative regulator of the root hair formation that determines the pattern of root hairs on the root epidermis of *Arabidopsis thaliana* [17]. An investigation of this MYB based on its crystal structure revealed the DNA binding site AACNGC and also suggests that this MYB is able to differentiate between DNA methylation states [20]. DEFECTIVE IN TAPETAL DEVELOPMENT AND FUNCTION/MYB35 determines the sex of *Asparagus officinalis*, but not in all other species of this genus [21]. The *A. thaliana* PFG1/MYB12 and the paralogs PFG2/MYB11 and PFG3/MYB111 are responsible for the regulation of the flavonol biosynthesis in most tissues [18]. This process appears highly conserved as orthologs of the AtMYB11/AtMYB12/AtMYB111 clade in *Beta vulgaris* [22] and *Medicago truncatula* [19] are also regulators of the flavonol biosynthesis. MYB21 and MYB24 were identified as additional flavonol regulators in the stamen of *A. thaliana* [23]. Some processes like the regulation of the flavonol biosynthesis depend only on MYB regulators [18]. Other specialized metabolite biosynthesis pathways are regulated by the interaction of multiple proteins. The MBW complex, named after the three components MYB, bHLH, and WD40, is one of the best studied transcriptional regulation systems [24–27]. Two branches of the flavonoid biosynthesis, the anthocyanin and proanthocyanidin biosynthesis, are controlled by the MBW complex [24, 25, 28]. Since anthocyanins are responsible for the pigmentation of flowers and other plant structures, mutants in their regulation can be identified based on a visible phenotype. Proanthocyanidins are responsible for the coloration of seed coats thus mutants in their biosynthesis can be identified based on a yellow seed color. Mutations in the regulating MYBs and other transcription factors [29–32] are often the reason for loss of anthocyanins and/or proanthocyanins. Transcriptional regulation was studied based on these pathways due to their visually detectable phenotypes.

Since MYBs are controlling many processes in plants, there is also a substantial interest to understand their functions in crop species. In Brassicaceae, the glucosinolate content controlled by several MYBs is an economically relevant trait [33]. ATR1/MYB34, HIG1/MYB51, and MYB122 increase the indolic glucosinolates and HAG1/MYB28, PMG2/RAO7/MYB29, and MYB76 the aliphatic glucosinolates in *Arabidopsis thaliana* [34–36]. The red coloration of sugar beets is controlled by BvMYB1 which activates two betalain biosynthesis genes [37]. *AmMYB1* in amaranth was identified as best candidate gene to explain the seed coloration variation between accessions [38]. A MYB appears to be the underlying factor of post-harvest hardening that renders a specific yam accession inedible within a day [39, 40]. MYB duplications between different apple cultivars appear responsible for differences in the red fruit flesh coloration [41]. Consumers prefer apricots with a red blush which is controlled by an anthocyanin biosynthesis activating MYB [42]. The identification of MYB candidate genes and the regulated processes is the first step towards modification through SMART breeding or genome editing [43, 44]. This interest in MYBs sparked numerous genome-wide investigations in species with a new genome or transcriptome assembly [3, 4, 22, 31, 45–48]. The identification of MYBs is repeatedly performed on many different data sets with strong variation in the quality of the analyses. Well described *A. thaliana* MYB sequences [2] are often used as baits to find new MYBs based on sequence similarity. Like all routine tasks with clearly defined steps, the identification of MYBs is a promising target for an automatic approach. We previously developed an automatic workflow, called KIPEs, for the annotation of core flavonoid biosynthesis genes which could also be expanded to the annotation of transcription factor gene families [4]. However, KIPEs is optimized for the identification and assessment of enzymes based on conserved amino acids in the active center. One underlying assumption is a small number of gene copies per species, which is violated by the very large MYB gene family. Also it is technically possible to run KIPEs for the identification of a gene family, the performance decreases with gene family size. Many previous studies relied only on BLAST or added additional filters for the presence of conserved R2R3-MYB domains in candidate sequences [4, 22, 49]. The inspection of MYB domains is laborious when performed manually, but suitable to define a set of fully functional R2R3-MYBs. While specificity of this filtering approach is high, it suffers from a low sensitivity i.e. neglects degenerated copies which might have experienced neofunctionalization. There are other solutions to identify orthologous sequences in a large number of species independent of the presence of specific sequence patterns [50, 51], but these approaches would require a substantial amount of manual cleaning to narrow down a final set of MYB sequences. Particular challenges are analyses based on transcriptome assemblies, because transcriptome assemblies show often a large number of isoforms resulting from alternative splicing or artifacts [52, 53].

This study presents a Python-based pipeline to provide a high quality annotation of all MYBs in a given set of peptide or coding sequences that are provided as input. Additionally, MYB candidates are checked for conserved domains and assigned to orthologs in other plant species. Genome sequencing and the construction of assemblies is becoming a routine task. The generation of high quality structural annotations is also advancing quickly with the aid of massive RNA-Seq data sets and full length transcript sequencing. Therefore, a huge number of data sets will be available to study MYBs in an unprecedented number of different plant species. Our automatic identification of MYBs in a large number of species facilitates pan-MYB analyses to better understand the evolution of the MYBome and to transfer functional insights acquired in one species effectively to orthologs in other species.

## Implementation

### MYB data sets

The identification of MYBs in novel genome or transcriptome sequences requires broad phylogenetic coverage of bait sequences. The MYB domain sequences of *Arabidopsis thaliana* [2], *Vitis vinifera* [3], *Beta vulgaris* [22], *Musa acuminata* [4], *Vitis vinifera [3], Medicago truncatula, Populus trichocarpa, Citrus sinensis, Solanum lycopersicum, Solanum tuberosum, Aquilegia coerulea, Oryza sativa, Zea mays [47], Amborella trichopoda, Picea abies, Selaginella moellendorfii, Physcomitrella patens, Chlamydomonas reinhardtii, Volvox carteri, Micromonas pusilla, Ostreococcus lucimarinus, and Cyanidioschyzon merolae* [10] were merged to generate a bait sequence collection. Closely related non-MYB sequences including CDC5 were identified by running a BLASTp search with the bait MYB sequences against the Araport11 peptide sequences [54]. Hits with a minimum BLAST score of 100 were collected and stripped of any *bona fide* MYBs. This step allowed the identification of MYB-like sequences, but excludes spurious hits that would slow down the following analysis steps. While the previously described MYBs form a collection of 1889 ingroup sequences, these 26 non-MYB sequences represent the outgroup sequences for down-stream analysis.

### Pipeline

The automatic annotation pipeline is summarized in **Fig. 1**. Required inputs are (1) the MYB bait sequences (described above), (2) a classification of the MYB bait sequences into ingroup and outgroup, and (3) a set of coding sequences or peptide sequences that will be analyzed. Step 0: All parameters, tool versions, and the input files are logged in a report file for reproducibility. If the Python module hashlib is available, md5sums are calculated for all input files to ensure an accurate documentation. However, calculation of this checksum is optional and file names (including their paths) will be documented in any case. Cleaning of the input sequences removes any characters from the sequence names that would interfere with the phylogenetic analysis. Step 1: Initial candidates are identified based on local sequence similarity via BLAST [55, 56] or HMMER [57]. Default parameters accept BLASTp hits with 50 amino acid length, 80% alignment similarity, and a maximum of 100 hits per bait sequence. This is a very sensitive setting given the large number of bait sequences and an expected MYB gene family size below 300 in most species. If a collection of coding sequences is provided, these will be translated and then compared based on BLASTp to harness the stronger conservation of amino acid sequences compared to nucleotide sequences. Step 3: A phylogenetic tree is constructed with these initial candidates and all bait sequences. Alignments are constructed with MAFFT [58]. Tree construction via RAxML [59] and FastTree2 [60] is supported. FastTree2 (-wag -nopr -nosupport) is recommended due to substantially higher speed when large data sets are analyzed. Step 3: Calculation of distances between different leaves of the tree is used to identify ortholog relationships between bait and candidate sequences. The Python package dendropy [61] is applied using the patristic distance method and counting edges. To exclude outliers caused by fragmented sequences or annotation artifacts, candidates are excluded if the distance to the next bait sequences exceeds three times the average distance of nearest neighbours. This cutoff was optimized by manually inspecting distributions of this value in context with the corresponding phylogenetic trees, but it is possible to modify this value as well as most other parameters. The bait sequences with the shortest distance are identified for each selected candidate in the tree. If most of these bait sequences are ingroup MYBs, the candidate is classified as MYB (Additional file 1). If most of these bait sequences are outgroup MYBs, the candidate is classified as a MYB-like sequence (aka non-MYB). All sequences passing this filter are considered clean candidates. Step 4: A check for the presence of MYB repeats is performed based on regular expressions (see documentation for details) derived from previously reported alignments [10, 62]. A repeat-based MYB classification is widely used and also supported here. However, it is important to note that these groups do not represent monophyletic groups. Step 5: A new phylogenetic tree is constructed with the clean MYB candidates and all bait sequences. Step 6: An optional step assigns all newly discovered MYBs to a group of reference MYBs e.g. the well characterized *A. thaliana* MYBs. Based on the assumptions that orthologs are likely to have the same functions, this generates hypotheses about the function of the newly discovered MYBs. Additionally, it is possible to identify the expansion and contraction of specific MYB lineages compared to this reference. Step 7: It is possible to collapse large groups of very similar sequences in the analyzed data set and to represent these by only the longest sequence. This option is intended for transcriptome assemblies which can include large numbers of isoforms caused by alternative splicing and artifacts. Step 8: A new phylogentic tree of the representative sequences identified in step 7 and the reference sequence set (e.g. *A. thaliana* MYBs) is constructed.

**Fig. 1:**
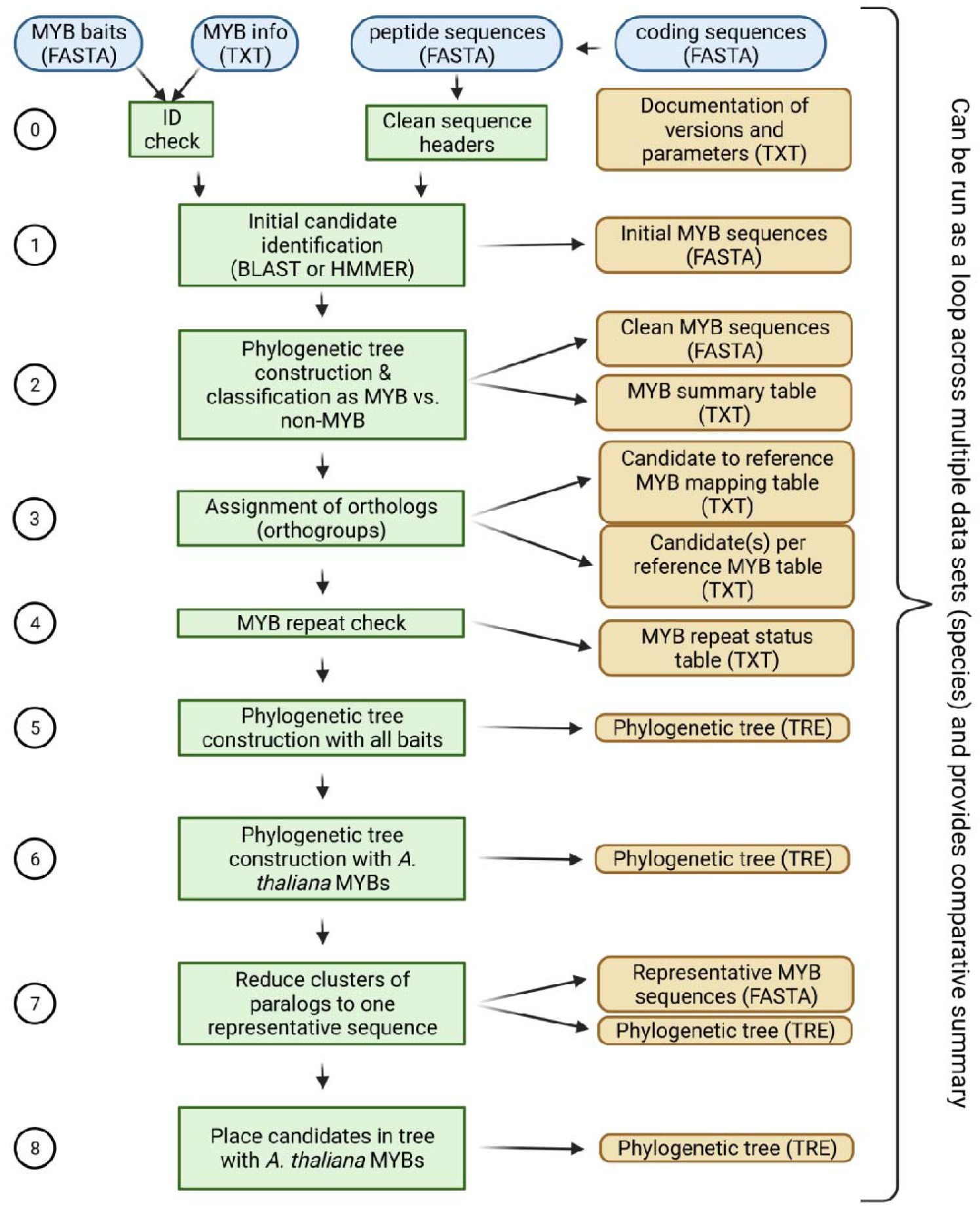
Simplified illustration of a pipeline for the automatic annotation of MYBs. Please refer to the text and the documentation on github for additional details about the pipeline. There is an option to run this pipeline across all provided input files in a folder. This enables the generation of summary files that compare the MYB gene families between the analyzed species.

## Results and Discussion

### Proof of concept and benchmarking

Several benchmarking data sets were analyzed to ensure that the pipeline performs well for a range of different plant species. Araport11 sequences of the *A. thaliana* accession Col-0 [54] were analyzed and the well characterized MYBs were recovered (example output files on github). This demonstrates that the pipeline works as expected. The annotated sequences of the *A. thaliana* accession Nd-1 [63] were screened for MYBs. As expected, there is a 1:1 relationship between the MYBs of Col-0 and Nd-1 (Additional file 2). This demonstrates that not just identical, but also slightly different sequences are accurately identified.

### Performance

Benchmarking and performance tests were performed on a compute cluster without control over other jobs running on the same machine. This prevented a precise and informative calculation of run times, but also represents realistic conditions. A total of 121 coding sequence sets were downloaded from Phytozome [64] and screened for MYBs. The average run time per species using default parameters, 4 CPUs, FastTree, and v0.153 of the pipeline was about 8 minutes (**Fig. 2**, Additional file 3). The memory requirements of all steps in the pipeline are very low (<1GB). The major factor contributing to the run time is the construction of a phylogenetic tree. However, the use of RAxML takes substantially longer. If a job is canceled, the analysis can continue at the last completed check point or at the last analyzed data set (species), respectively. The required hard disk space is minimal (63.5 MB for *A. thaliana*). Changes in the parameters and especially in the number of supplied bait sequences can alter the computational costs substantially. While there are differences with respect to the run time depending on the number of predicted species per family and the size of the MYB gene family, this analysis indicates that large data sets can be processed effectively.

**Fig. 2:**
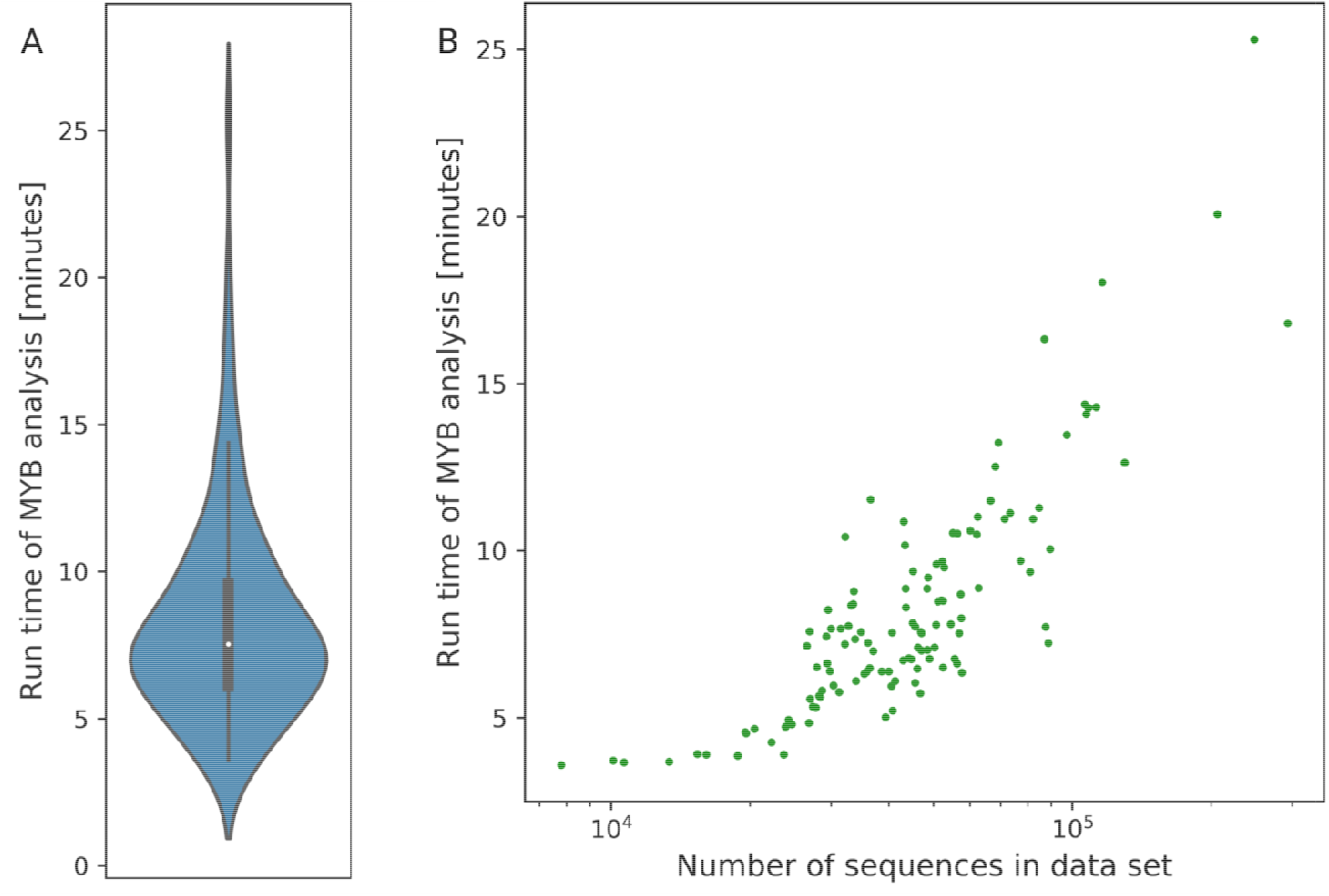
Average run time per data set (species) with default parameters based on coding sequences (A). Positive correlation of run time with the number of sequences in the data set (B).

### Discovery of MYBs in the *Castanea crenata* genome sequence

As a proof of concept, MYBs encoded in the recently sequenced *Castanea crenata* genome [65] were investigated. The predicted peptide sequences were screened with default parameters and resulted in the identification of 136 MYBs (**Fig. 3**, Additional file 4). A R2R3-MYB domain was detected in 112 of them. No orthologs of the *Cruciferae*-specific glucosinolate biosynthesis regulating MYBs AtMYB028, AtMYB029, AtMYB034, AtMYB051, AtMYB076, and AtMYB122 were detected in *C. crenata*. This is not surprising, because *C. crenata* belongs to the *Fagaceae*, and also in line with previous reports about the absence of this MYB linage from non-Cruciferae [3]. Regulators of the flavonoid biosynthesis (AtMYB011/AtMYB012/AtMYB111, Ccr1.0Bg1101.1-S7/Ccr1.0Jg2696.1-S7), anthocyanin biosynthesis (AtMYB075/AtMYB090/AtMYB113/AtMYB114, Ccr1.0Ag5288.1-S6), and proanthocyanidin biosynthesis (AtMYB123, Ccr1.0Ag1758.1 / Ccr1.0Ag1766.1 / Ccr1.0Ag1768.1 / Ccr1.0Ag1770.1 / Ccr1.0Ag1773.1 / Ccr1.0Ag5531.1 / Ccr1.0Ag5542.1 / Ccr1.0Ag5543.1 / Ccr1.0Eg0443.1 / Ccr1.0Eg0444.1 / Ccr1.0Gg0097.1 / Ccr1.0Gg0098.1 / Ccr1.0Hg2677.1 / Ccr1.0Jg0953.1 / Ccr1.0Jg2532.1 / Ccr1.0Lg0385.1 / Ccr1.0Lg3370.1) were detected. In general, copy number differences can be explained by lineage-specific duplication events. Therefore, it is not possible to establish 1:1 relationships between *A. thaliana* and *C. crenata*. Interestingly, there are multiple homologs of AtMYB123 in *C. crenata* which could indicate a high importance of proanthocyanidins in this species. Further investigations could analyze the transcription of these genes to exclude unexpressed copies. The reduced number of PAP homologs could suggest a lower importance of anthoycanins. While the copies of the anthocyanin regulator show different functions in *A. thaliana* [66], loss of MYB114 in several *A. thaliana* accessions including Col-0 [67] and Nd-1 [68] suggest that there is functional redundancy between them.

**Fig. 3:**
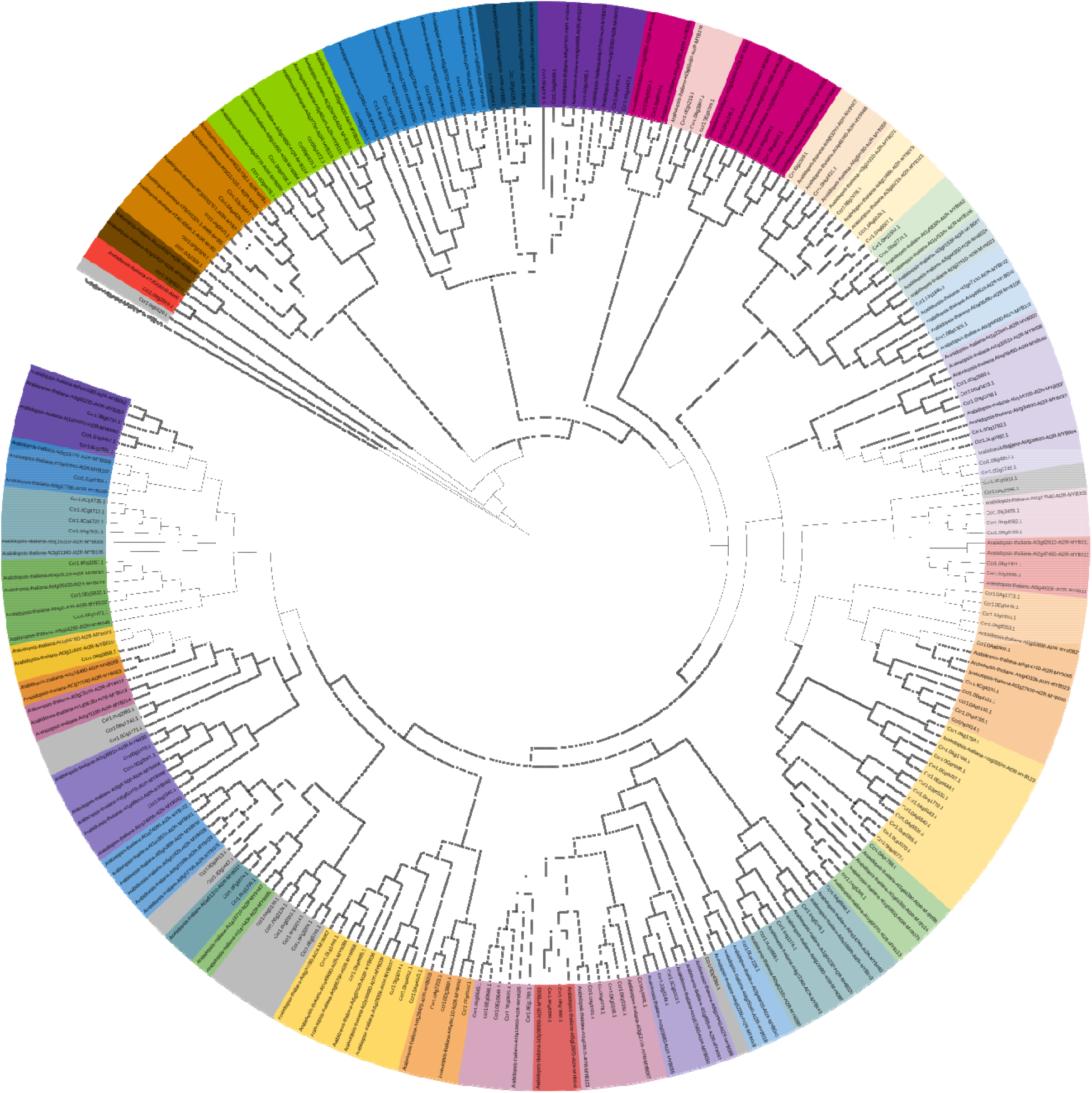
Relationships of *Castanea crenata* MYB candidates and well characterized *Arabidopsis thaliana* MYBs. This figure was constructed with iTOL [69].

### Discovery of MYBs in the Croton tiglium transcriptome assembly

To demonstrate that the pipeline also works for inherently incomplete transcriptome assemblies, MYBs were investigated in the transcriptome assembly of *Croton tiglium* [70]. An analysis with default parameters revealed 140 MYBs (**Fig. 4**, Additional file 5). This includes 103 candidates with a complete R2R3-MYB domain. Transcriptome assemblies are known to be rich in isoforms of the same genes, which can be due to alternative splicing or artifacts. Clusters of these isoforms were represented by the longest sequence among them. This reduced the number of MYB candidates to 79 including 69 with a R2R3 domain. Not all *A. thaliana* MYBs are matched by orthologs in *C. tiglium*. Although this transcriptome assembly is based on paired-end RNA-seq data sets representing leaf, root, stem, seed, and inflorescence, some not or lowly expressed MYBs might not be represented in the assembly. Therefore, they cannot be identified in this analysis. Again, the absence of orthologs of the glucosinolate regulating MYBs aligns well with previous reports [3], because *C. tiglium* belongs to the Euphorbiaceae. The conserved regulators of the flavonoid biosynthesis (AtMYB011/AtMYB012/AtMYB111, TRINITY-DN21046-c0-g1-i2-S7) and proanthocyanidin biosynthesis (AtMYB123, TRINITY-DN31260-c4-g2-i2-S5) were detected. These findings are in line with previous reports of the flavonol regulators and proanthocyanidin regulator being detectable in this transcriptome assembly [71]. The absence of a PAP ortholog from the assembly is not surprising, because none of the sampled tissues showed a pigmentation by anthocyanins [70]. Anthocyanin regulators are well known to be lowly expressed in tissues without anthocyanin pigmentation [72, 73] thus a lack of expression is a likely explanation of this result.

**Fig. 4:**
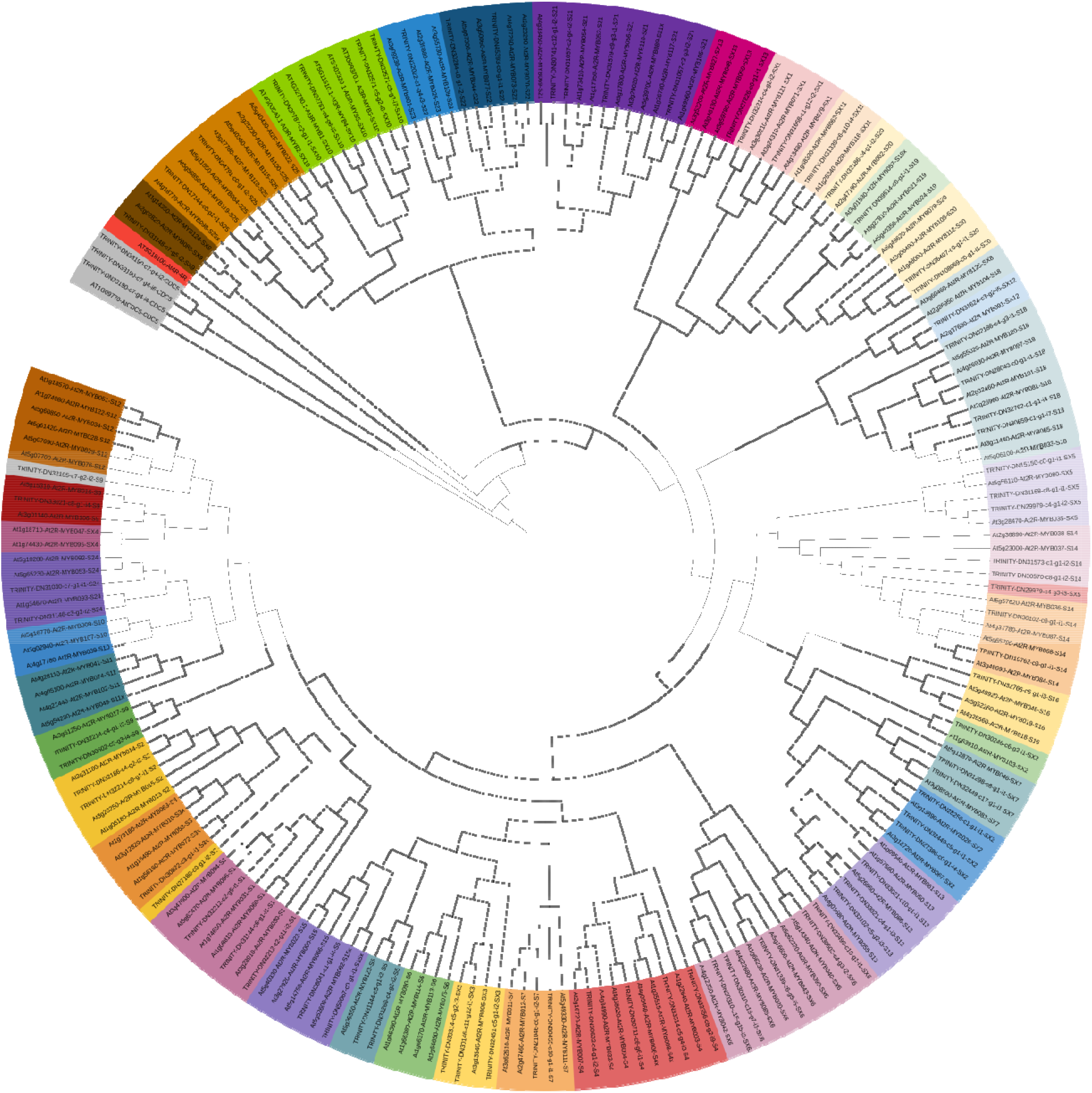
Relationships of *Croton tiglium* and well characterized *Arabidopsis thaliana* MYBs. This figure was constructed with iTOL [69].

### Limitations and next steps

The collection of bait sequences distributed with the tool should be appropriate for most applications. This collection covers a large taxonomic range including chlorophytes, charophycean algae, bryophytes, and vascular plants. The dominance of sequences belonging to vascular plants can be explained by the generally larger MYB families in these species. The major MYB lineages are represented in this collection, but MYB lineages that are restricted to certain taxonomic groups could be missing. While the initial identification based on overall sequence similarity is a robust approach, the precise classification and functional annotation of such MYBs could be less accurate due to the lack of close orthologs. The only critical step is the separation between MYBs and MYB-like sequences. Emerging and species-specific MYB clades embedded in widely distributed clades are not a major concern, because it is possible to adjust the parameters of the analysis when analyzing isolated species e.g. members of the Lycophytes or Magnoliidae (see documentation for technical details). Only sequences at the basis of the MYB gene family tree could be at risk of being missed.

There are constrains that influence the size of the bait sequence collection. A narrower set of bait sequences could reduce the run time if a comprehensive investigation of numerous genome sequences of one taxonomic group is planned. An analysis with a more comprehensive set of MYB sequences could improve the quality of the results, but would also substantially increase the run time of the analysis. The generation of an ideal bait sequence set which represents the complete phylogenetic diversity of MYBs with a minimal number of sequences is a task for future studies.

Most steps are deterministic, but minor variations might occur as part of the tree building. However, no biologically relevant differences were observed during the analyses of 121 benchmarking datasets. Additionally, the results of analyses with BLAST-based selection of candidates were consistent with the results of corresponding analyses using HMMER for the identification of initial candidates.

## Conclusions

This approach only relies on standard tools which should be installed on most systems and are also easy to install if not available already. Technical checks on *A. thaliana* datasets indicate that the pipeline is accurately identifying MYBs. The performance allows the investigation of one species within minutes on ubiquitously available hardware. An investigation of the MYB gene families in *Castanea crenata* and *Croton tiglium* revealed expected patterns and demonstrated the potential to analyze transcriptome and genome sequence assemblies. While this approach is dedicated to the analysis of MYBs, it could be adjusted to investigate other transcription factor gene families.

## Availability and requirements

**Project name:** MYB_annotator

**Project home page:** https://github.com/bpucker/MYB_annotator

**Operating system(s):** Linux

**Programming language:** Python3

**Other requirements:** dendropy, BLAST, HMMER, MAFFT, RAxML or FastTree2

**License:** GNU General Public License v3.0

**Any restrictions to use by non-academics:** none

## Declarations

### Ethics approval and consent to participate

Not applicable

### Consent for publication

Not applicable

### Availability of data and materials

The pipeline and the required input data sets are freely available via github: https://github.com/bpucker/MYB_annotator. Additionally, the released version was archived via zenodo (10.5281/zenodo.6174039). The analyzed transcriptomic and genomic resources are publicly available: *Castanea crenata* (BPMU01000001-BPMU01000781) and *Croton tiglium* (PRJNA416498).

### Authors’ contributions

BP planned the study, wrote the software, performed the bioinformatic analysis, interpreted the results, and wrote the manuscript.

## Acknowledgements

Many thanks to the German network for bioinformatics infrastructure (de.NBI, grant 031A533A) and the Bioinformatics Resource Facility (BRF) at the Center for Biotechnology (CeBiTec) at Bielefeld University for providing an environment to perform the computational analyses. I am also grateful to Hanna Marie Schilbert for testing some earlier versions, providing feedback, and requesting additional features. bioRender.com was used to construct some of the figures.

## References

1. Klempnauer KH, Gonda TJ, Bishop JM. Nucleotide sequence of the retroviral leukemia gene v-myb and its cellular progenitor c-myb: the architecture of a transduced oncogene. Cell. 1982;31 2 Pt 1:453–63.

2. Stracke R, Werber M, Weisshaar B. The R2R3-MYB gene family in Arabidopsis thaliana. Current Opinion in Plant Biology. 2001;4:447–56.

3. Matus JT, Aquea F, Arce-Johnson P. Analysis of the grape MYB R2R3 subfamily reveals expanded wine quality-related clades and conserved gene structure organization across Vitis and Arabidopsis genomes. BMC Plant Biology. 2008;8:83.

4. Pucker B, Pandey A, Weisshaar B, Stracke R. The R2R3-MYB gene family in banana (Musa acuminata): Genome-wide identification, classification and expression patterns. PLOS ONE. 2020;15:e0239275.

5. Jiang C-K, Rao G-Y. Insights into the Diversification and Evolution of R2R3-MYB Transcription Factors in Plants. Plant Physiology. 2020;183:637–55.

6. Yuan Y, Yang X, Feng M, Ding H, Khan MT, Zhang J, et al. Genome-wide analysis of R2R3-MYB transcription factors family in the autopolyploid Saccharum spontaneum: an exploration of dominance expression and stress response. BMC Genomics. 2021;22:622.

7. Dubos C, Stracke R, Grotewold E, Weisshaar B, Martin C, Lepiniec L. MYB transcription factors in Arabidopsis. Trends in Plant Science. 2010;15:573–81.

8. Ogata K, Kanei-Ishii C, Sasaki M, Hatanaka H, Nagadoi A, Enari M, et al. The cavity in the hydrophobic core of Myb DNA-binding domain is reserved for DNA recognition and trans-activation. Nat Struct Mol Biol. 1996;3:178–87.

9. Jia L, Clegg MT, Jiang T. Evolutionary dynamics of the DNA-binding domains in putative R2R3-MYB genes identified from rice subspecies indica and japonica genomes. Plant Physiol. 2004;134:575–85.

10. Du H, Liang Z, Zhao S, Nan M-G, Tran L-SP, Lu K, et al. The Evolutionary History of R2R3-MYB Proteins Across 50 Eukaryotes: New Insights Into Subfamily Classification and Expansion. Sci Rep. 2015;5:11037.

11. Ito M. Conservation and diversification of three-repeat Myb transcription factors in plants. J Plant Res. 2005;118:61–9.

12. Haga N, Kato K, Murase M, Araki S, Kubo M, Demura T, et al. R1R2R3-Myb proteins positively regulate cytokinesis through activation of KNOLLE transcription in Arabidopsis thaliana. Development. 2007;134:1101–10.

13. Rosinski JA, Atchley WR. Molecular evolution of the Myb family of transcription factors: evidence for polyphyletic origin. J Mol Evol. 1998;46:74–83.

14. Braun EL, Grotewold E. Newly discovered plant c-myb-like genes rewrite the evolution of the plant myb gene family. Plant Physiol. 1999;121:21–4.

15. Kranz H, Scholz K, Weisshaar B. c-MYB oncogene-like genes encoding three MYB repeats occur in all major plant lineages. Plant J. 2000;21:231–5.

16. Jiang C, Gu J, Chopra S, Gu X, Peterson T. Ordered origin of the typical two-and three-repeat Myb genes. Gene. 2004;326:13–22.

17. Lee MM, Schiefelbein J. WEREWOLF, a MYB-Related Protein in Arabidopsis, Is a Position-Dependent Regulator of Epidermal Cell Patterning. Cell. 1999;99:473–83.

18. Stracke R, Ishihara H, Huep G, Barsch A, Mehrtens F, Niehaus K, et al. Differential regulation of closely related R2R3-MYB transcription factors controls flavonol accumulation in different parts of the Arabidopsis thaliana seedling. Plant J. 2007;50:660–77.

19. Naik J, Rajput R, Pucker B, Stracke R, Pandey A. The R2R3-MYB transcription factor MtMYB134 orchestrates flavonol biosynthesis in Medicago truncatula. Plant Mol Biol. 2021;106:157–72.

20. Wang B, Luo Q, Li Y, Yin L, Zhou N, Li X, et al. Structural insights into target DNA recognition by R2R3-MYB transcription factors. Nucleic Acids Research. 2020;48:460–71.

21. Harkess A, Huang K, van der Hulst R, Tissen B, Caplan JL, Koppula A, et al. Sex Determination by Two Y-Linked Genes in Garden Asparagus. The Plant Cell. 2020;32:1790–6.

22. Stracke R, Holtgräwe D, Schneider J, Pucker B, Rosleff Sörensen T, Weisshaar B. Genome-wide identification and characterisation of R2R3-MYB genes in sugar beet (Beta vulgaris). BMC Plant Biology. 2014;14:249.

23. Zhang X, He Y, Li L, Liu H, Hong G. Involvement of the R2R3-MYB transcription factor MYB21 and its homologs in regulating flavonol accumulation in Arabidopsis stamen. Journal of Experimental Botany. 2021;72:4319–32.

24. Walker AR, Davison PA, Bolognesi-Winfield AC, James CM, Srinivasan N, Blundell TL, et al. The TRANSPARENT TESTA GLABRA1 locus, which regulates trichome differentiation and anthocyanin biosynthesis in Arabidopsis, encodes a WD40 repeat protein. Plant Cell. 1999;11:1337–50.

25. Nesi N, Debeaujon I, Jond C, Pelletier G, Caboche M, Lepiniec L. The TT8 Gene Encodes a Basic Helix-Loop-Helix Domain Protein Required for Expression of DFR and BAN Genes in Arabidopsis Siliques. Plant Cell. 2000;12:1863–78.

26. Ramsay NA, Glover BJ. MYB-bHLH-WD40 protein complex and the evolution of cellular diversity. Trends Plant Sci. 2005;10:63–70.

27. Lloyd A, Brockman A, Aguirre L, Campbell A, Bean A, Cantero A, et al. Advances in the MYB–bHLH–WD Repeat (MBW) Pigment Regulatory Model: Addition of a WRKY Factor and Co-option of an Anthocyanin MYB for Betalain Regulation. Plant and Cell Physiology. 2017;58:1431–41.

28. Baudry A, Heim MA, Dubreucq B, Caboche M, Weisshaar B, Lepiniec L. TT2, TT8, and TTG1 synergistically specify the expression of BANYULS and proanthocyanidin biosynthesis in Arabidopsis thaliana. Plant J. 2004;39:366–80.

29. Nesi N, Jond C, Debeaujon I, Caboche M, Lepiniec L. The Arabidopsis TT2 Gene Encodes an R2R3 MYB Domain Protein That Acts as a Key Determinant for Proanthocyanidin Accumulation in Developing Seed. Plant Cell. 2001;13:2099–114.

30. Jin W, Wang H, Li M, Wang J, Yang Y, Zhang X, et al. The R2R3 MYB transcription factor PavMYB10.1 involves in anthocyanin biosynthesis and determines fruit skin colour in sweet cherry (Prunus avium L.). Plant Biotechnology Journal. 2016;14:2120–33.

31. Wang H, Zhang H, Yang Y, Li M, Zhang Y, Liu J, et al. The control of red colour by a family of MYB transcription factors in octoploid strawberry (Fragaria × ananassa) fruits. Plant Biotechnology Journal. 2020;18:1169–84.

32. Zheng X, Om K, Stanton KA, Thomas D, Cheng PA, Eggert A, et al. The regulatory network for petal anthocyanin pigmentation is shaped by the MYB5a/NEGAN transcription factor in Mimulus. Genetics. 2021;217.

33. Tan Z, Xie Z, Dai L, Zhang Y, Hu Z, Tang S, et al. Genome-and transcriptome-wide association studies reveal the genetic basis and the breeding history of seed glucosinolate content in Brassica napus. Plant Biotechnol J. 2021. https://doi.org/10.1111/pbi.13707.

34. Gigolashvili T, Berger B, Mock H-P, Müller C, Weisshaar B, Flügge U-I. The transcription factor HIG1/MYB51 regulates indolic glucosinolate biosynthesis in Arabidopsis thaliana. The Plant Journal. 2007;50:886–901.

35. Hirai MY, Sugiyama K, Sawada Y, Tohge T, Obayashi T, Suzuki A, et al. Omics-based identification of Arabidopsis Myb transcription factors regulating aliphatic glucosinolate biosynthesis. PNAS. 2007;104:6478–83.

36. Frerigmann H, Gigolashvili T. Update on the role of R2R3-MYBs in the regulation of glucosinolates upon sulfur deficiency. Frontiers in Plant Science. 2014;5:626.

37. Hatlestad GJ, Akhavan NA, Sunnadeniya RM, Elam L, Cargile S, Hembd A, et al. The beet Y locus encodes an anthocyanin MYB-like protein that activates the betalain red pigment pathway. Nat Genet. 2015;47:92–6.

38. Stetter MG, Vidal-Villarejo M, Schmid KJ. Parallel Seed Color Adaptation during Multiple Domestication Attempts of an Ancient New World Grain. Molecular Biology and Evolution. 2020;37:1407–19.

39. Siadjeu C, Pucker B, Viehöver P, Albach DC, Weisshaar B. High Contiguity de novo Genome Sequence Assembly of Trifoliate Yam (Dioscorea dumetorum) Using Long Read Sequencing. Genes. 2020;11:274.

40. Siadjeu C, Mayland-Quellhorst E, Pande S, Laubinger S, Albach DC. Transcriptome Sequence Reveals Candidate Genes Involving in the Post-Harvest Hardening of Trifoliate Yam Dioscorea dumetorum. Plants. 2021;10:787.

41. Chagné D, Lin-Wang K, Espley RV, Volz RK, How NM, Rouse S, et al. An Ancient Duplication of Apple MYB Transcription Factors Is Responsible for Novel Red Fruit-Flesh Phenotypes. Plant Physiol. 2013;161:225–39.

42. Xi W, Feng J, Liu Y, Zhang S, Zhao G. The R2R3-MYB transcription factor PaMYB10 is involved in anthocyanin biosynthesis in apricots and determines red blushed skin. BMC Plant Biology. 2019;19:287.

43. Mikhaylova EV, Shein MY, Artyukhin AY, Sukhareva AS, Panfilova MA, Kuluev BR. Editing of the MYB genes in Brassica napus as a method to increase anthocyanin pigmentation and stress tolerance. E3S Web Conf. 2020;224:04022.

44. Khusnutdinov E, Sukhareva A, Panfilova M, Mikhaylova E. Anthocyanin Biosynthesis Genes as Model Genes for Genome Editing in Plants. International Journal of Molecular Sciences. 2021;22:8752.

45. Yanhui C, Xiaoyuan Y, Kun H, Meihua L, Jigang L, Zhaofeng G, et al. The MYB Transcription Factor Superfamily of Arabidopsis: Expression Analysis and Phylogenetic Comparison with the Rice MYB Family. Plant Mol Biol. 2006;60:107–24.

46. Wilkins O, Nahal H, Foong J, Provart NJ, Campbell MM. Expansion and Diversification of the Populus R2R3-MYB Family of Transcription Factors. Plant Physiology. 2009;149:981–93.

47. Du H, Feng B-R, Yang S-S, Huang Y-B, Tang Y-X. The R2R3-MYB Transcription Factor Gene Family in Maize. PLOS ONE. 2012;7:e37463.

48. Cao Y, Jia H, Xing M, Jin R, Grierson D, Gao Z, et al. Genome-Wide Analysis of MYB Gene Family in Chinese Bayberry (Morella rubra) and Identification of Members Regulating Flavonoid Biosynthesis. Frontiers in Plant Science. 2021;12:1244.

49. Li Z, Peng R, Tian Y, Han H, Xu J, Yao Q. Genome-Wide Identification and Analysis of the MYB Transcription Factor Superfamily in Solanum lycopersicum. Plant and Cell Physiology. 2016;57:1657–77.

50. Yang Y, Moore MJ, Brockington SF, Soltis DE, Wong GK-S, Carpenter EJ, et al. Dissecting Molecular Evolution in the Highly Diverse Plant Clade Caryophyllales Using Transcriptome Sequencing. Mol Biol Evol. 2015;32:2001–14.

51. Emms DM, Kelly S. OrthoFinder: phylogenetic orthology inference for comparative genomics. Genome Biology. 2019;20:238.

52. Zhang R, Kuo R, Coulter M, Calixto CPG, Entizne JC, Guo W, et al. A high resolution single molecule sequencing-based Arabidopsis transcriptome using novel methods of Iso-seq analysis. 2021.

53. Guang A, Howison M, Zapata F, Lawrence C, Dunn CW. Revising transcriptome assemblies with phylogenetic information. PLOS ONE. 2021;16:e0244202.

54. Cheng C-Y, Krishnakumar V, Chan AP, Thibaud-Nissen F, Schobel S, Town CD. Araport11: a complete reannotation of the Arabidopsis thaliana reference genome. The Plant Journal. 2017;89:789–804.

55. Altschul SF, Gish W, Miller W, Myers EW, Lipman DJ. Basic local alignment search tool. J Mol Biol. 1990;215:403–10.

56. Altschul SF, Madden TL, Schäffer AA, Zhang J, Zhang Z, Miller W, et al. Gapped BLAST and PSI-BLAST: a new generation of protein database search programs. Nucleic Acids Res. 1997;25:3389–402.

57. Mistry J, Finn RD, Eddy SR, Bateman A, Punta M. Challenges in homology search: HMMER3 and convergent evolution of coiled-coil regions. Nucleic Acids Res. 2013;41:e121.

58. Katoh K, Standley DM. MAFFT Multiple Sequence Alignment Software Version 7: Improvements in Performance and Usability. Mol Biol Evol. 2013;30:772–80.

59. Kozlov AM, Darriba D, Flouri T, Morel B, Stamatakis A. RAxML-NG: a fast, scalable and user-friendly tool for maximum likelihood phylogenetic inference. Bioinformatics. 2019;35:4453–5.

60. Price MN, Dehal PS, Arkin AP. FastTree 2 – Approximately Maximum-Likelihood Trees for Large Alignments. PLOS ONE. 2010;5:e9490.

61. Sukumaran J, Holder MT. DendroPy: a Python library for phylogenetic computing. Bioinformatics. 2010;26:1569–71.

62. Feng G, Burleigh JG, Braun EL, Mei W, Barbazuk WB. Evolution of the 3R-MYB Gene Family in Plants. Genome Biology and Evolution. 2017;9:1013–29.

63. Pucker B, Holtgräwe D, Stadermann KB, Frey K, Huettel B, Reinhardt R, et al. A chromosome-level sequence assembly reveals the structure of the Arabidopsis thaliana Nd-1 genome and its gene set. PLOS ONE. 2019;14:e0216233.

64. Goodstein DM, Shu S, Howson R, Neupane R, Hayes RD, Fazo J, et al. Phytozome: a comparative platform for green plant genomics. Nucleic Acids Res. 2012;40 Database issue:D1178–86.

65. Shirasawa K, Nishio S, Terakami S, Botta R, Marinoni DT, Isobe S. Chromosome-level genome assembly of Japanese chestnut (Castanea crenata Sieb. et Zucc.) reveals conserved chromosomal segments in woody rosids. 2021.

66. Koo Y, Poethig RS. Expression pattern analysis of three R2R3-MYB transcription factors for the production of anthocyanin in different vegetative stages of Arabidopsis leaves. Applied Biological Chemistry. 2021;64:5.

67. Gonzalez A, Zhao M, Leavitt JM, Lloyd AM. Regulation of the anthocyanin biosynthetic pathway by the TTG1/bHLH/Myb transcriptional complex in Arabidopsis seedlings. The Plant Journal. 2008;53:814–27.

68. Pucker B, Holtgräwe D, Sörensen TR, Stracke R, Viehöver P, Weisshaar B. A De Novo Genome Sequence Assembly of the Arabidopsis thaliana Accession Niederzenz-1 Displays Presence/Absence Variation and Strong Synteny. PLOS ONE. 2016;11:e0164321.

69. Letunic I, Bork P. Interactive Tree Of Life (iTOL) v5: an online tool for phylogenetic tree display and annotation. Nucleic Acids Research. 2021;49:W293–6.

70. Haak M, Vinke S, Keller W, Droste J, Rückert C, Kalinowski J, et al. High Quality de Novo Transcriptome Assembly of Croton tiglium. Front Mol Biosci. 2018;5.

71. Pucker B, Reiher F, Schilbert HM. Automatic Identification of Players in the Flavonoid Biosynthesis with Application on the Biomedicinal Plant Croton tiglium. Plants. 2020;9:1103.

72. Takos AM, Jaffé FW, Jacob SR, Bogs J, Robinson SP, Walker AR. Light-Induced Expression of a MYB Gene Regulates Anthocyanin Biosynthesis in Red Apples. Plant Physiol. 2006;142:1216–32.

73. Guo N, Han S, Zong M, Wang G, Zheng S, Liu F. Identification and differential expression analysis of anthocyanin biosynthetic genes in leaf color variants of ornamental kale. BMC Genomics. 2019;20:564.

