## Supplementary figures and images for "Automatic identification and annotation of MYB gene family members in plants"

### Additional file 1

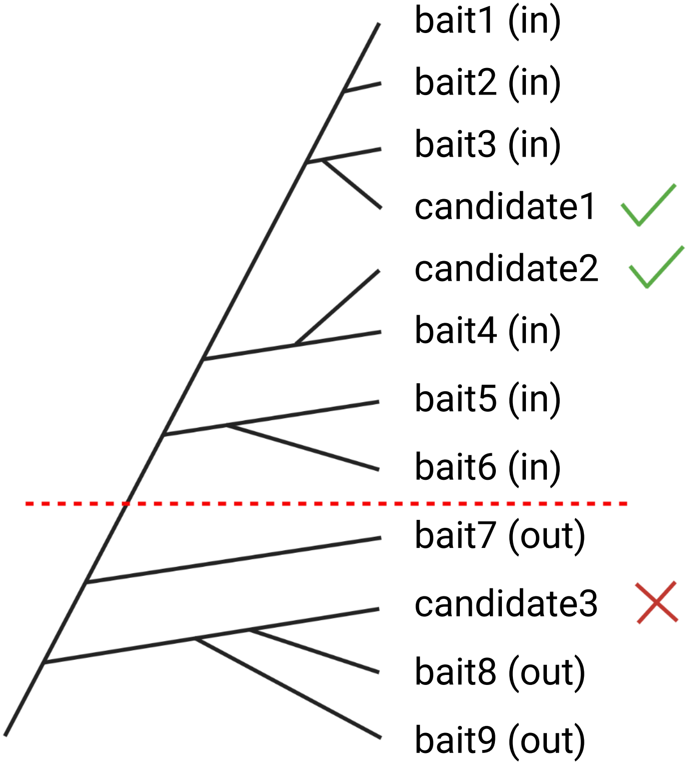
